# Computational and experimental optimization of T cell activation

**DOI:** 10.1101/629857

**Authors:** Bulent Arman Aksoy, Eric Czech, Chrystal Paulos, Jeff Hammerbacher

**Affiliations:** Microbiology and Immunology Department at the Medical University of South Carolina

## Abstract

Bead-based activation is widely-used for *ex vivo* expansion of T cells for either research or clinical purposes. Despite its wide use, culture conditions that can potentially affect the efficiency of bead-based T cell activation has not been extensively documented. With the help of computationally-driven experimental investigations of basic culturing factors, we found that culture density, bead-to-cell ratio, and debeading time can have a major impact on the efficiency of bead-based T cell activation for short-term cultures. Furthermore, discrepancies across expected and observed activation efficiencies helped discover interesting artifacts of bead-based T cell activation.

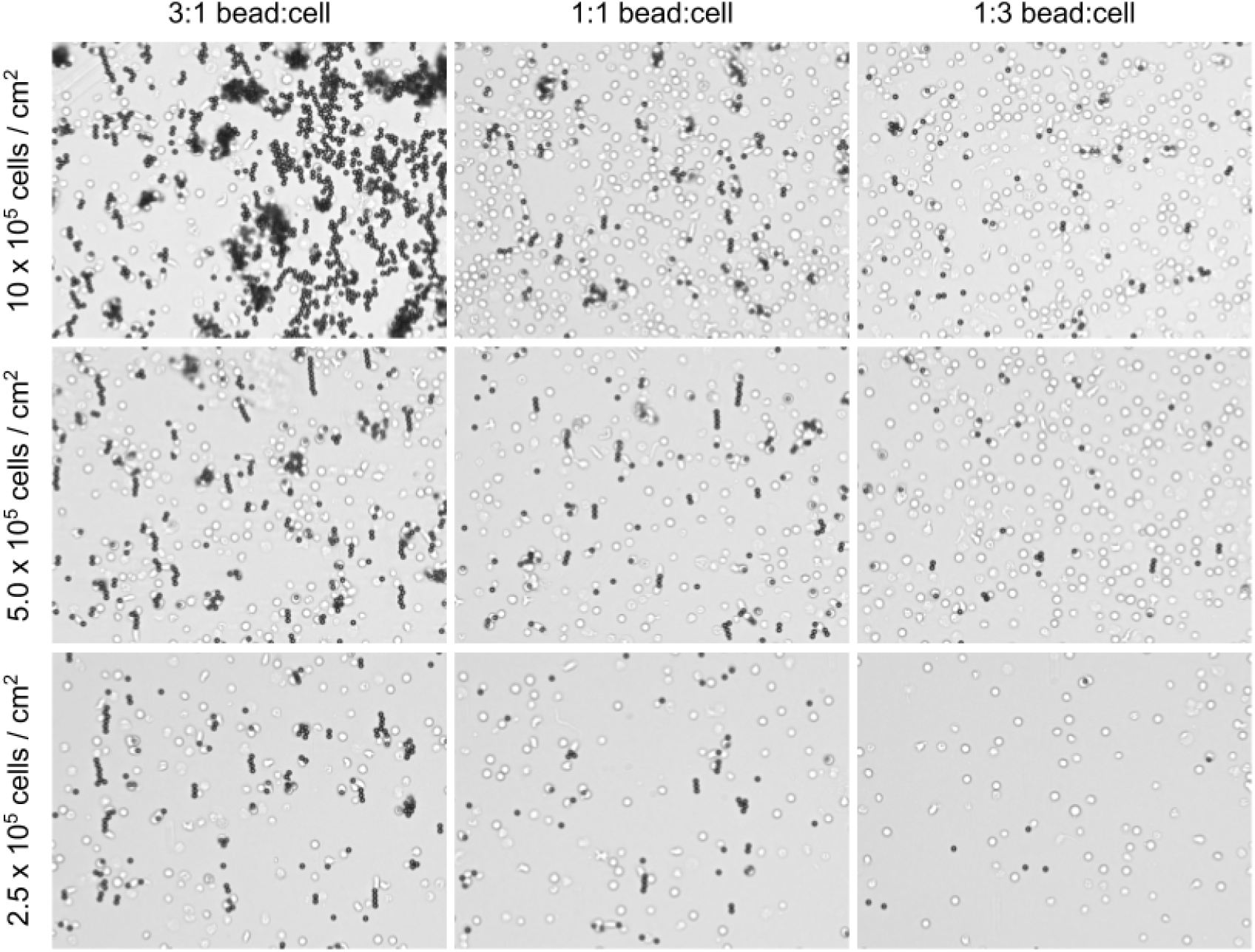
Human primary T cells were imaged together with activation beads at 20X magnification after three hours of culturing at varying confluencies and bead-to-cell ratios.

---

Many laboratories use CD3/CD28-activator magnetic beads for short- and long-term expansion of human primary T cells for research or clinical purposes (Trickett and Kwan 2003). Without activation, T cells do not proliferate efficiently and furthermore, their metabolically and epigenetically inactive profile makes them harder to manipulate. Different activation techniques can favor different T cell subtypes and therefore, the choice of the activation technique can be made based on the specific use case or experimental setup (Li and Kurlander 2010). Activation with anti-CD3 and anti-CD28 coated beads is preferred in cases where the polyclonal nature of the T cell population needs to preserved, for example T cell expansion for adoptive cell therapy for cancer or infectious disease. Even though the bead-based activation is widely-used and well-known, the factors that might affect the overall efficiency of the cellular product are poorly documented (Kalamasz et al. 2004).

Because bead-based activation requires physical contact between the cell and the bead, it has long been assumed that it is important to supply an unstimulated T cell population with enough beads to insure all cells are contacting at least one bead. The recommended bead-to-cell ratio is 1-to-1 for <u>Dynabeads™ Human T-Activator CD3/CD28 for T Cell Expansion and Activation</u>. Given that each bead can bind to multiple cells and each cell can bind to multiple beads, we asked if we could use fewer than recommended beads and still activate all the cells. We have previously showed that it is possible to titrate the number of beads down to 2-4 fold without compromising the activation efficiency (Aksoy et al., n.d.); but we have not investigated the potential impact of other cofactors, such as the confluency–the fraction of physical space occupied by cells and beads within a given culture vessel.

Unstimulated primary T cells do not adhere to the culture dish and therefore, they are virtually immobile. Likewise, magnetic beads are also immobile and due their density, similar to unstimulated T cells, they do not float. Hence we assume that when the culture is being established, the cells and the beads will randomly be distributed throughout the bottom of the vessel. We also propose that cells that are next to a bead will get activated, and cells will remain unstimulated if they are not touching a bead. These assumptions allow us to computationally model the initial activation process through simulations and explore the parameters that can impact the efficiency of activation. For simplicity, we defined our activation efficiency as the fraction of cells that got activated in a simulated environment and we explored the expected outcome of varying the bead-to-cell ratio and the confluency across 100 simulations per condition. We found that the activation efficiency was dependent on both the confluency and the bead-to-cell ratio, where as the confluency got lower, the activation efficiency also got lower at a given bead-to-cell ratio. This means that optimization of bead-to-cell ratio should not be generalized across different cultures unless the seeding confluency is fixed. For example, our simulations showed that working with a 3-to-1 cell-to-bead ratio at 12.5% confluency and 1-to-9 ratio at 100% confluency both yielded similar activation efficiencies: 0.53∓0.06 and 0.52∓0.01, respectively (**Figure 1**).

**Figure 1:**
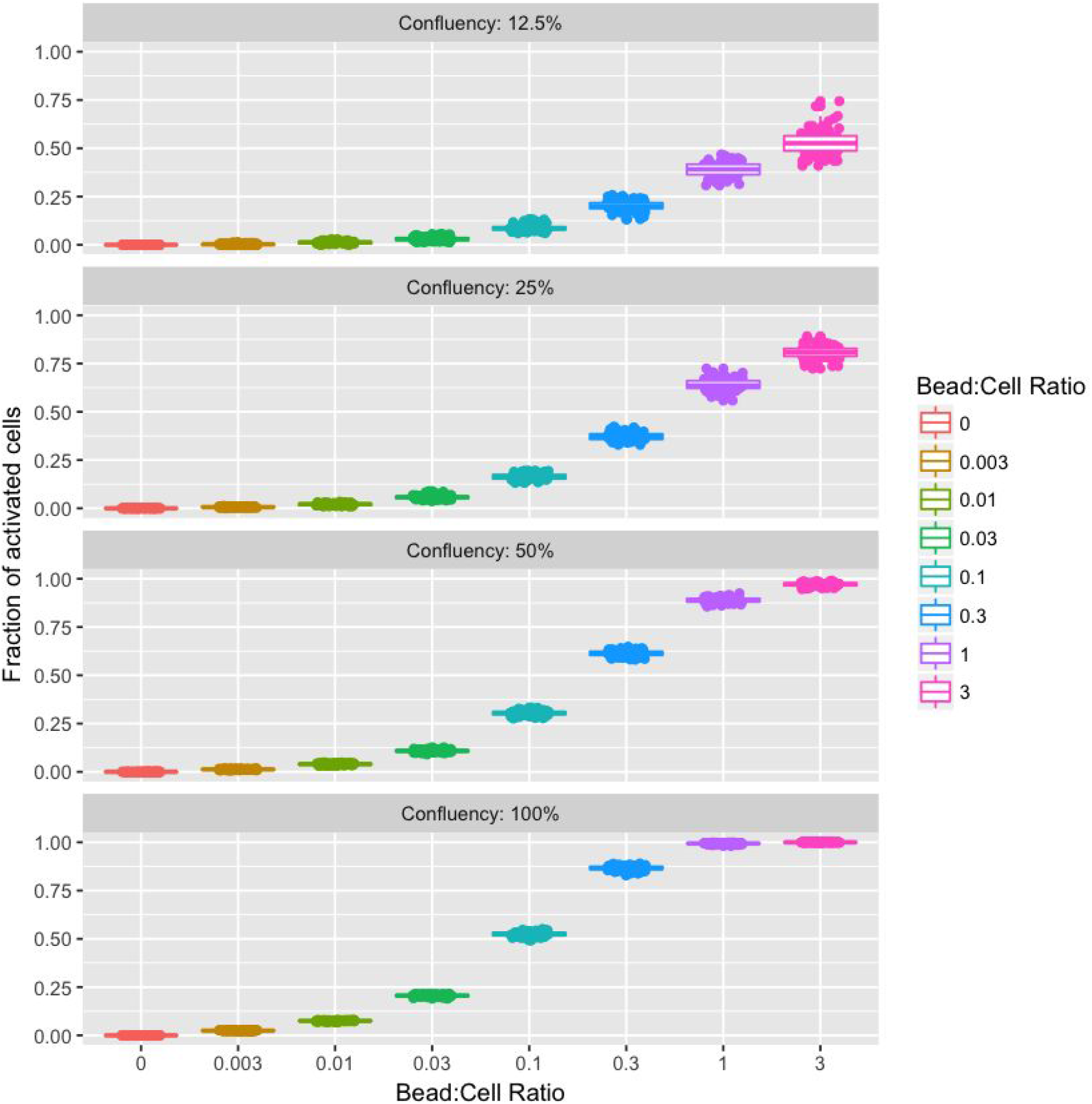
Simulations of T cell cultures that are being activated with aCD3/aCD28 beads help predict expected activation efficiency (fraction of activated cells) based on culture confluency and bead-to-cell ratio. The simulations showed that the higher the confluency, the better the activation efficiency became at a given bead-to-cell ratio. Therefore, at relatively higher confluencies, the activation efficiency did not immediately suffer when the beads were titrated down–*e.g*. activating a culture at a 1-to-3 cell-to-bead ratio rather than the recommended-to-1 ratio (source: <u>Modeling bead-based T cell activation on a population level</u>).

When cultured with or without IL-2 supplement in standard T-cell media, unstimulated cells can survive for at least 4 weeks but the longer they are in the culture, the worse the overall survival gets (see **Supplemental Figure 1a**). Although naive cells (CCR7+CD45RO-) disappear from the unstimulated population relatively faster, they are predominantly alive within the first week (see **Supplemental Figure 1b**). Given that bead-based activation requires physical contact, naive cells that do not contact any beads during the culture should stay as naive and naive cells that do contact at least one bead should get activated and differentiate. Furthermore, it takes a naive cell (CCR7+CD45RO-) at least 3 days to turn into a central memory cell (CCR7+CD45RO+) upon bead activation (see **Supplemental Figure 2**). Therefore, we can experimentally estimate the activation efficiency across multiple conditions by counting the number of naive cells that are left in a culture after a 3-day activation. Our results showed that, when the confluency was kept constant (1 million cells per 2 cm^2^), decreasing the bead-to-cell ratio caused fewer naive cells to get activated (see **Figure 2**). Furthermore, our observed activation efficiencies roughly corresponded to the hypothetical 25% confluency (see **Figure 1**).

**Figure 2:**
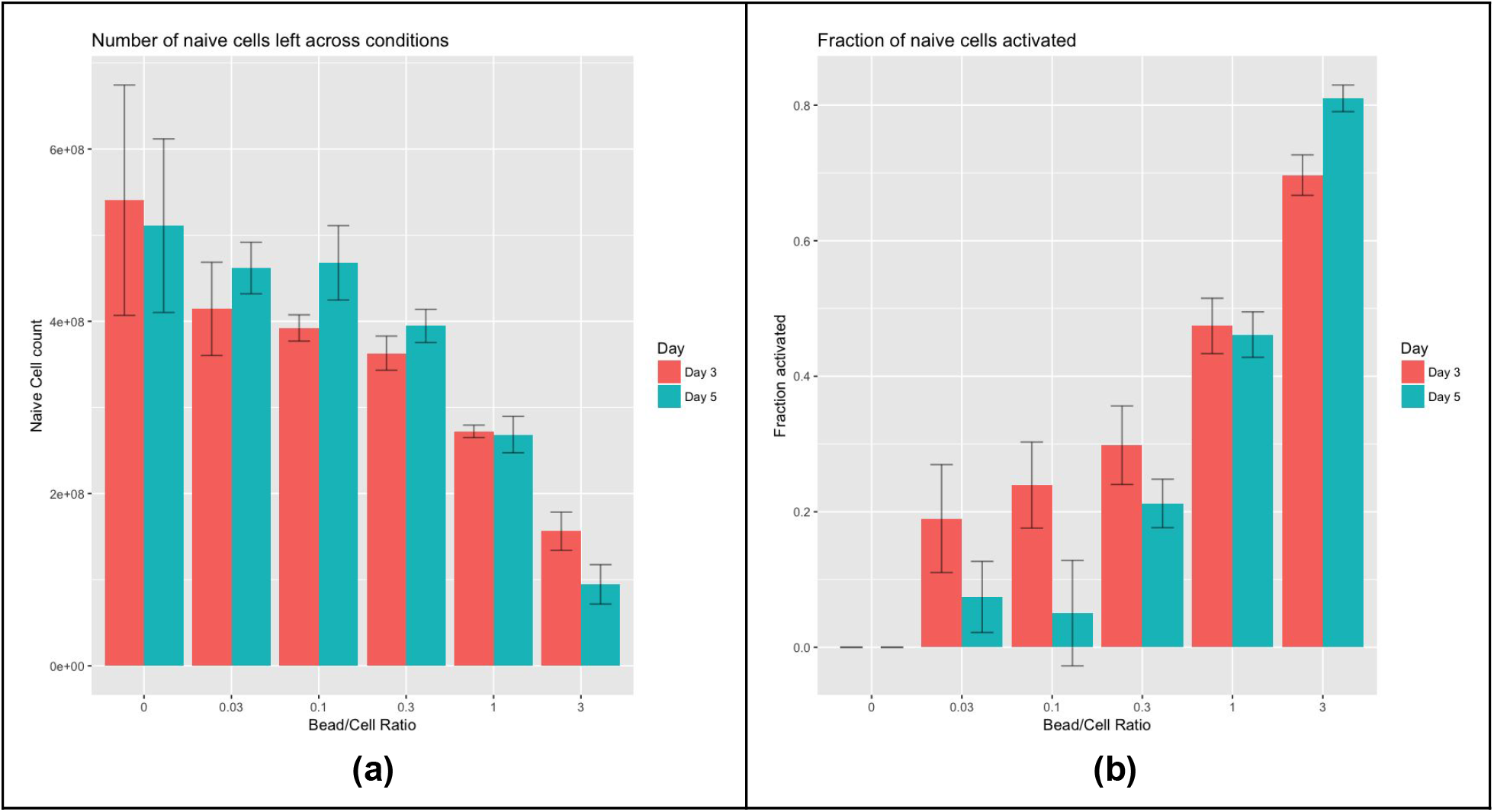
Use of fewer beads leads to activation of fewer naive T cells (CCR7+ CD45RO-) as predicted by the simulations. (a) When the same number of cells were activated using varying cell-to-bead ratios, the number of naive cells that are left in the culture (both on days 3 and 5) decreased as more beads were used. (b) When the number of naive cells left in the culture were normalized against the no-bead (i.e. no activation) condition, the fraction of naive cells left in the culture followed a trend that was similar to the outcome of the simulations, specifically the simulations at 25% confluency (source: <u>Titrate beads and count naive cells post-activation</u>).

We then wanted to experimentally verify our computational predictions about the effect of varying confluencies on the activation efficiency. At a fixed cell-to-bead ratio of 1-to-1, we tried to activate unstimulated T cells at varying concentrations and found the optimal confluency to be between 0.5 and 1.5 million cells per 2 cm^2^ (see **Figure 3** and **Supplemental Figure 3**). Given that this roughly corresponds to our hypothetical 25% confluency, we were surprised to see that the activation efficiency was relatively low at confluencies higher than 2 million cells per cm^2^ as our simulations were predicting higher efficiencies at higher confluencies. These data can potentially be explained by the fixed volume (1.5 mL per 2 cm^2^) of culture media (and hence the nutrients in it) starting to become limiting to the cell activation and proliferation. However, it is not straightforward to experimentally validate this without using specialized culture flasks since using higher volumes of media can easily start to have a negative impact on the gas exchange rate.

**Figure 3:**
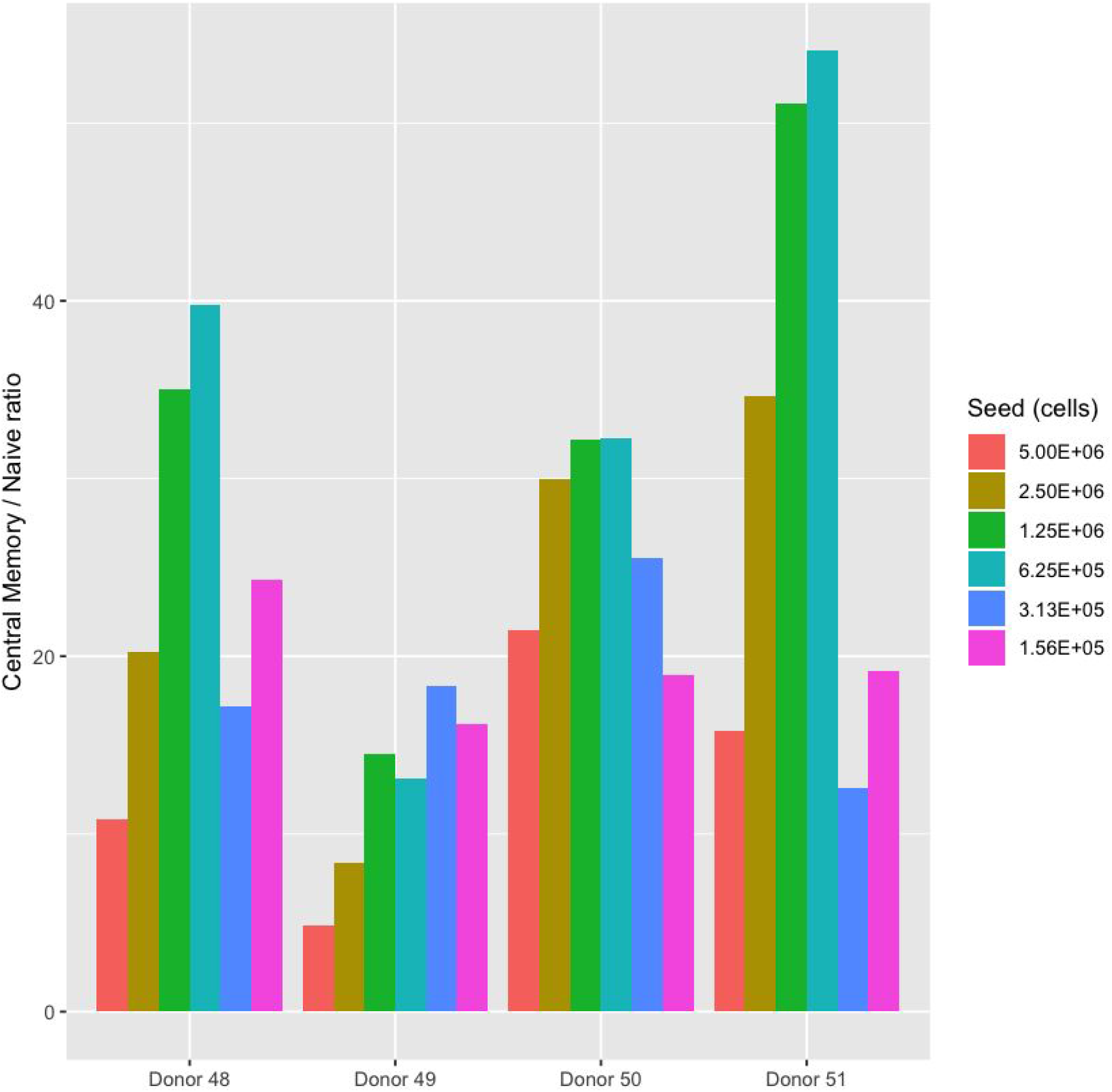
Seeding density can affect the activation efficiency. When we activated T cells at a recommended bead-to-cell ratio (1-to-1) but varied the amount of cells seeded in the culture, we saw that relatively high or low seeding densities led to worse activation efficiency. Given that the number of naive cells seeded varied across different densities, we estimated the activation efficiency through central memory (CM: CCR7+ CD45RO+) to naive (CCR7+ CD45RO-) ratio (**Supplemental Figure 3**). Cultures seeded with 625,00-1,250,000 cells (per 1 cm^2^) had relatively better activation efficiencies compared to other confluencies. Relatively low activation efficiencies seen at higher confluencies might be due to the nutrient limitations, an artifact of working at higher cell densities (source: <u>Titrate cells and beads to measure the effect of seeding density on activation efficiency</u>).

During our bead titration trials, we have consistently seen that using more than recommended beads yields relatively fewer central memory (CCR7+CD45RO+) cells (**Figure 4** and **Supplemental Figure 4**). This result was unexpected because we have previously concluded that using more beads led to a higher fraction of activated T cells. One critical part of the bead-based activation is the debeading process: once the cells are fully activated with beads (within 2-3 days), the beads are commonly removed from the culture as they can interfere with downstream applications (e.g. electroporation of cells). Although debeading process should free a majority of the activated cells from the beads, if use of more beads causes more cells to get stuck on the beads, the overall yield can suffer due to loss of bead-bound and recently-activated cells at the end of the activation. We found that we could distinguish beads, T cells, and T cells that are bound to beads with the help of flow cytometry since they have distinct forward and side scatter profiles (**Supplemental Figure 5**). This allowed us to estimate the number of cells that got stuck on beads (and hence removed) during the debeading process. We found that regardless of the day of debeading, the number of cells that got stuck on the beads increases as the bead-to-cell ratio increased (**Figure 5a**). Furthermore, the cells that got stuck on beads were predominantly the activated cells (CD69+ by the second day) that were transitioning into a memory state (CD45O+ by the third day) (**Supplemental Figure 6** and **Supplemental Figure 7**). If debeading cells upon activation causes considerable loss of recently-activated cells as we have seen, then debeading at early time points upon activation should also lead to fewer T cells with a central memory phenotype (CCR7+ CD45RO+). This is expected because these lost cells comprise a fraction of the population with high proliferative capacity and will not have the opportunity to divide following transition to the central memory phenotype. To estimate the effect of the debeading time on the final central memory yield (CCR7+ CD45RO+), we debeaded T cells that were being activated at a 1-to-1 cell-to-bead ratio at different time points and then compared the yield on the 6th day of the culture. We found that debeading after only 1 day of activation almost always resulted in relatively fewer central memory (CCR7+ CD45RO+) cells compared to cultures that were debeaded on day 2 or day 3 (**Figure 5b**). When cells were seeded at a high confluency (2 million cells per 2 cm^2^), the yields of the cultures that were debeaded on different days were comparable.

**Figure 4:**
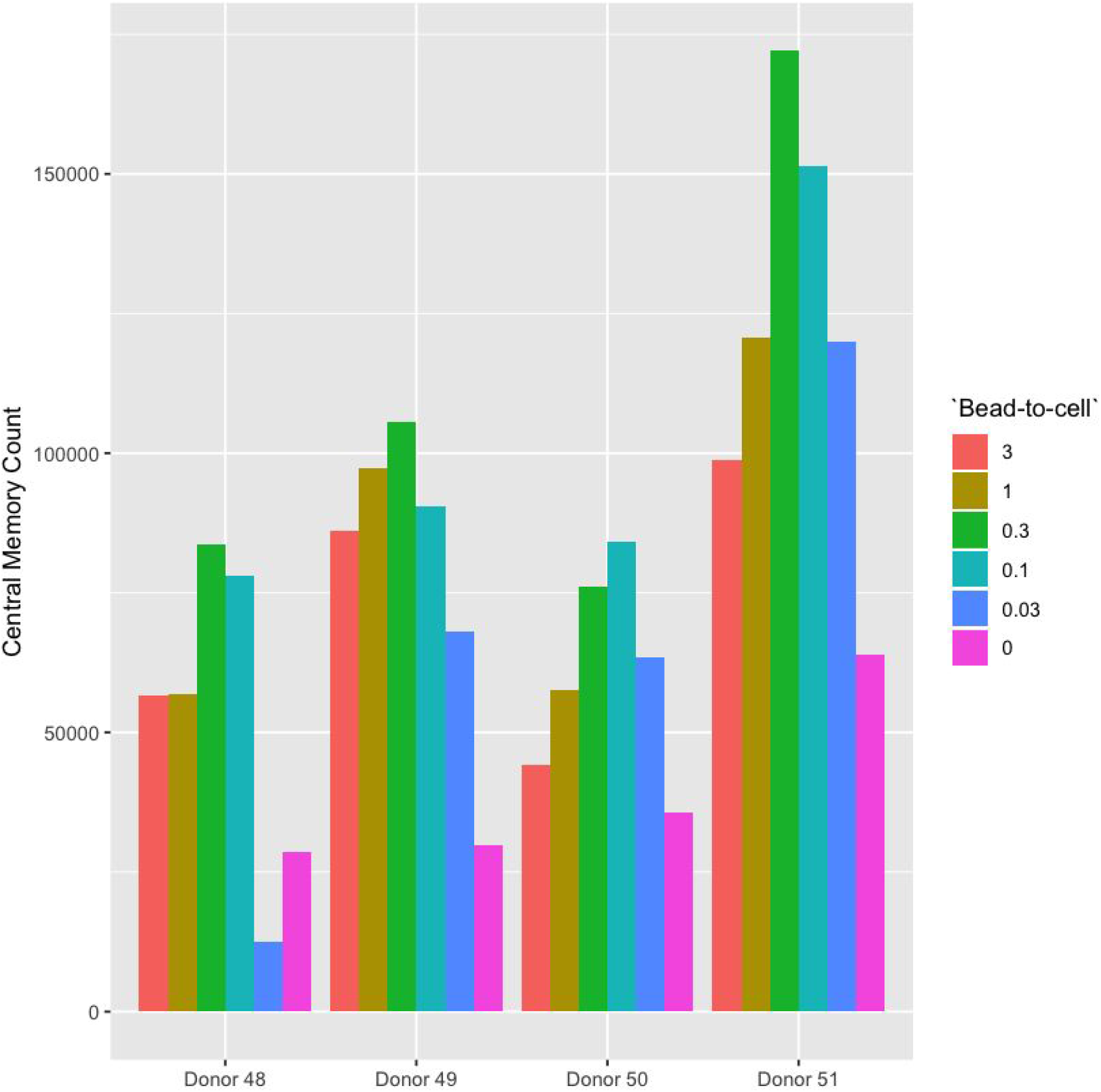
Surprisingly, activating T cells at relatively higher bead-to-cell ratios consistently yield fewer central memory (CM: CCR7+ CD45RO+) cells (**Supplemental Figure 4**). Although our simulations predicted that using more beads at a fixed confluency (25%) virtually resulted in higher activation efficiencies, our observations were in disagreement with this prediction: the central memory yield was almost always better at 1-to-3 and 1-to-9 bead-to-cell ratios compared to relatively higher or lower ones on the third day of the culture (source: Estimate the number of cells stuck on beads)

**Figure 5:**
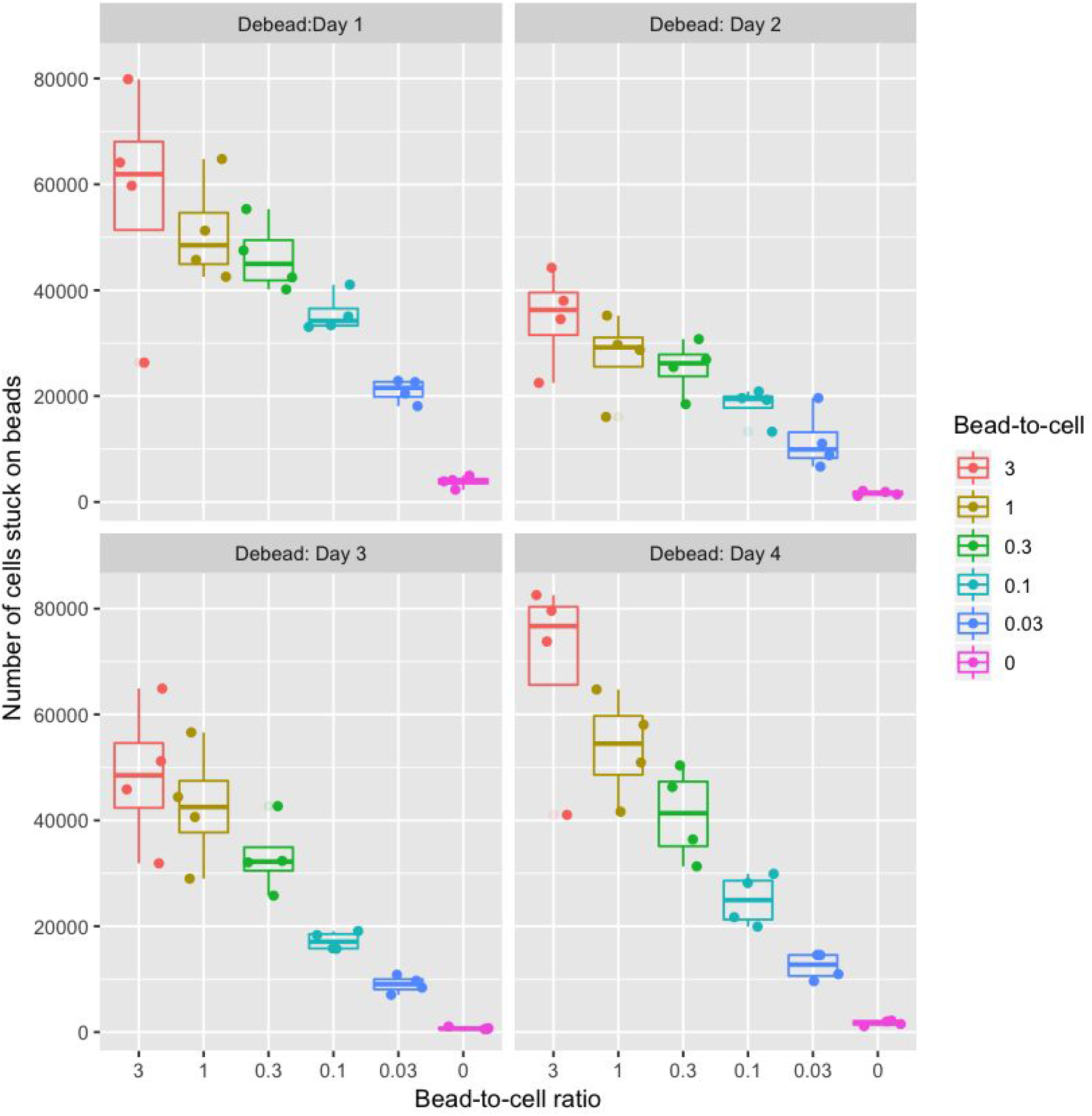
Use of more beads for activation causes more cells to get stuck on beads regardless of the duration of the culture, which might explain why we saw unexpected low central memory yields when T cell cultures were activated at more than 1-to-3 cell-to-bead ratios (source: <u>Estimate the number of cells stuck on beads</u>). Furthermore, we saw that these cells were predominantly activated and were transitioning into a central memory (CCR7+ CD45RO+) phenotype (**Supplemental Figures 6** and **7**).

**Figure 6:**
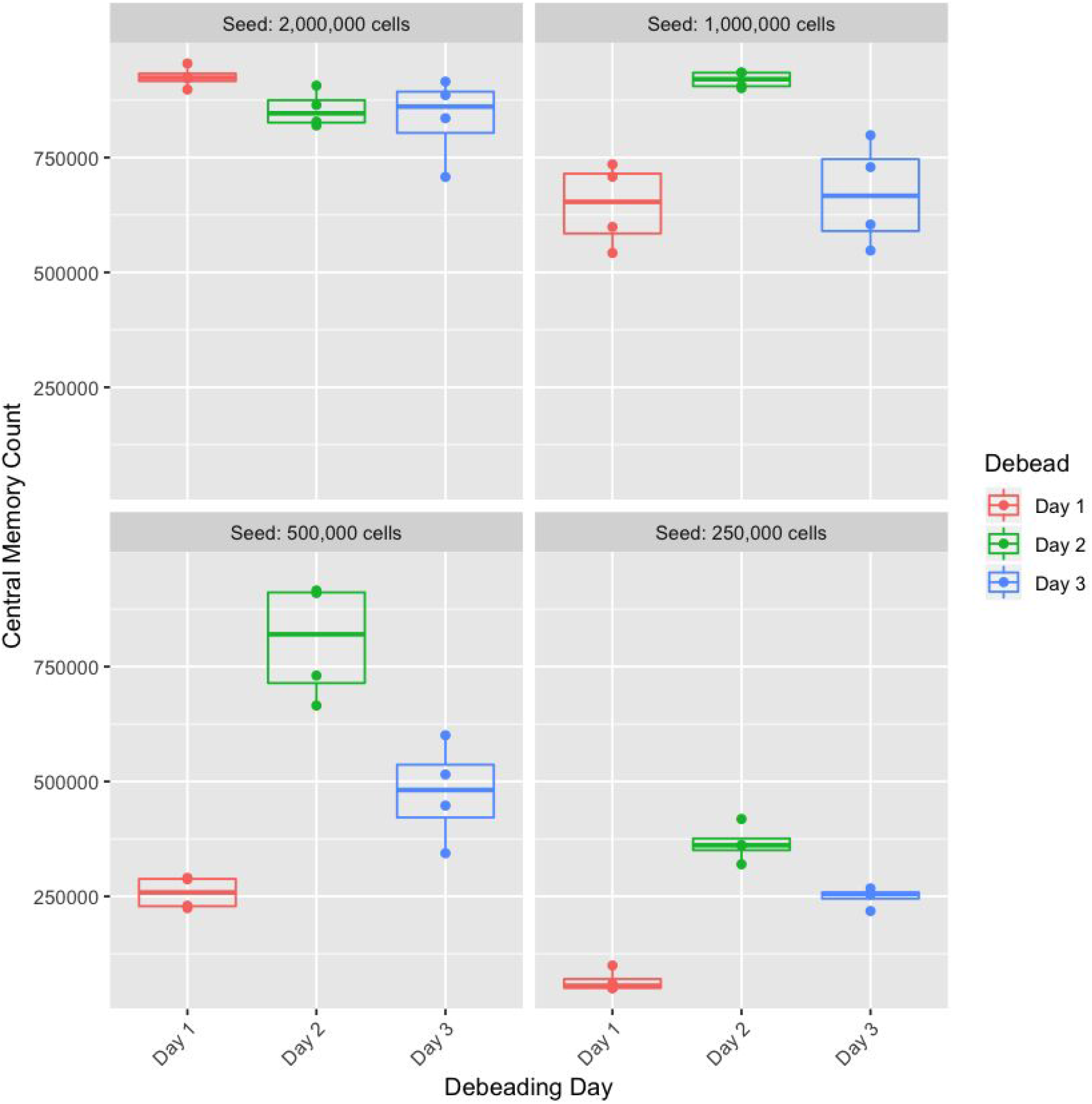
Debeading a T cell culture that is being activated with beads on day 1 (*i.e*. 24 hours after activation) almost always leads to lower central memory yield as measured on the 6th day of the culture. Except at 2 million cells per 2 cm^2^ density, debeading the culture on day 2 of the activation yielded more central memory (CM: CCR7+ CD45RO+) cells compared to debeading on day 3, which, in turn, yielded more compared to day 1 (source: Titrate cells and beads to measure the effect of seeding density on activation efficiency).

In summary, we show that modeling bead-based T cell activation as a physical process where the outcome is binary (a cell gets activated only if it is in direct contact with a bead) can help estimate the activation efficiency based on the culturing conditions. Experimental validation of these computational predictions further imply that seeding density, bead-to-cell ratio, and debeading time are all important factors that could impact the outcome of a bead-based T cell activation experiment. In our hands, activating T cells at a cell density of 0.5 to 1.5 million cells per 2 cm^2^ at a 1-to-9 to 1-to-1 bead-to-cell ratio and debeading on the second day led to optimal activation of the starting culture and yielded relatively more central memory cells. However, our results also show that the optimal activation setting is highly dependent on the particular culture settings used. Therefore, we highly encourage other scientists to optimize these parameters based on their unique experimental setup to achieve optimal activation efficiency for short-term T cell culturing experiments. Finally, we would like to emphasize that these considerations we have outlined in this manuscript are related to short-term T cell culturing experiments as slight differences in the activation efficiency are barely distinguishable for long term cultures (**Supplemental Figure 8**).

## Supplemental Materials and Methods

### Human primary T cell culture

Human primary T cells were isolated from healthy human buffy coats (purchased from Plasma Consultants LLC, Monroe Township, NJ) by EasySep™ Direct Human T Cell Isolation Kit (StemCell, Vancouver, Canada). Isolated T cells were initially frozen at 20 million per mL concentration in Thermo Fisher’s Recovery™ Cell Culture Freezing Medium (#12648010). When needed, the cells were thawed the day before the activation and kept in <u>T cell media (10.17504/protocols.io.qu5dwy6)</u>: RPMI with L-glutamine (Corning), 10% fetal bovine serum (Atlas Biologicals, Fort Collins, CO), 50 uM 2-mercaptoethanol (EMD Millipore), 25 mM HEPES (HyClone, GE Healthcare, Chicago, IL), 1% Penicillin-Streptomycin (Thermo Fisher), 1X sodium pyruvate (HyClone, GE Healthcare, Chicago, IL), and 1X non-essential amino acids (HyClone, GE Healthcare). T cells were activated with Dynabeads™ Human T-Activator CD3/CD28 for T Cell Expansion and Activation (Thermo Fisher, #11132D) within 24-well plates (with 2 cm^2^ surface area per well) and each sample was cultured within 1-1.5 mL of culture media without any IL-2 supplement. Unless otherwise noted, the working concentration was 1 million cells per mL.

### Data and Code Availability

All intermediate and final data sets together with their visualization logics are available at https://github.com/hammerlab/t-cell-activation. Specifically, the simulated T cell activation metrics are extracted from the following analysis: <u>Modeling bead-based T cell activation on a population level.ipynb</u>. In short, for each simulation, we allocated a 10×10 grid where each compartment can contain a T cell or a bead or it can be empty. Based on the virtual confluency (where 100% means we use all the compartments) and the bead-to-cell ratio (where 1:1 bead:cell ratio at 100% confluency means we place 50 beads and 50 T cells), we randomly place cells and beads within the grid. Once they are placed, we then assume a cell is activated only if at least one of its 8 neighboring compartments holds a bead. For each simulation, we estimate the efficiency through the fraction of T cells activated and for each condition, we pooled results from 100 independent simulations.

### Flow Cytometry

Flow cytometric analysis was performed on BD FACSVerse Flow Cytometer. Cells were collected and centrifuged at 300 x g for 5 minutes. The supernatant was aspirated. The cells were resuspended in flow buffer (PBS with 20% FBS) and the labeled-antibodies were added at the recommended concentration. The cells were stained at room temperature for 20-30 minutes at dark. After incubation, the cells were pelleted and resuspended in PBS. Flow cytometry results were analyzed by FlowJo v10 (TreeStar, Ashland, OR, USA).

### Antibodies

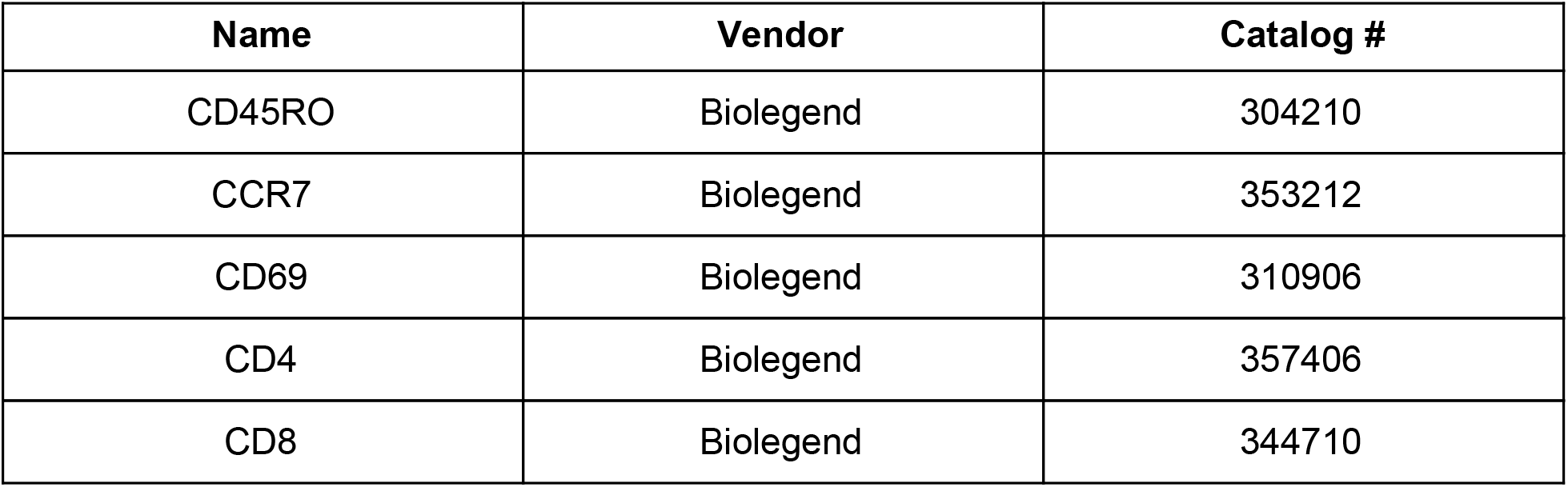

| **Name** | **Vendor** | **Catalog #** |
| --- | --- | --- |
| CD45RO | Biolegend | 304210 |
| CCR7 | Biolegend | 353212 |
| CD69 | Biolegend | 310906 |
| CD4 | Biolegend | 357406 |
| CD8 | Biolegend | 344710 |

## Acknowledgements

We thank Pinar Aksoy, Paulos Lab and Rubinstein Lab for their feedback on the manuscript and the project. This work is supported in part by the Flow Cytometry and Cell Sorting Unit Shared Resource, Hollings Cancer Center, Medical University of South Carolina (P30 CA138313). The authors declare no competing financial interests. We also would like to thank Google for supporting our work through providing research credits (#38362142) for the Google Cloud Platform. Finally, we would like to thank Nadia Eghbal for her generous donation towards our research project through a Helium Grant.

## Supplemental Figures

**Supplemental Figure 1:**
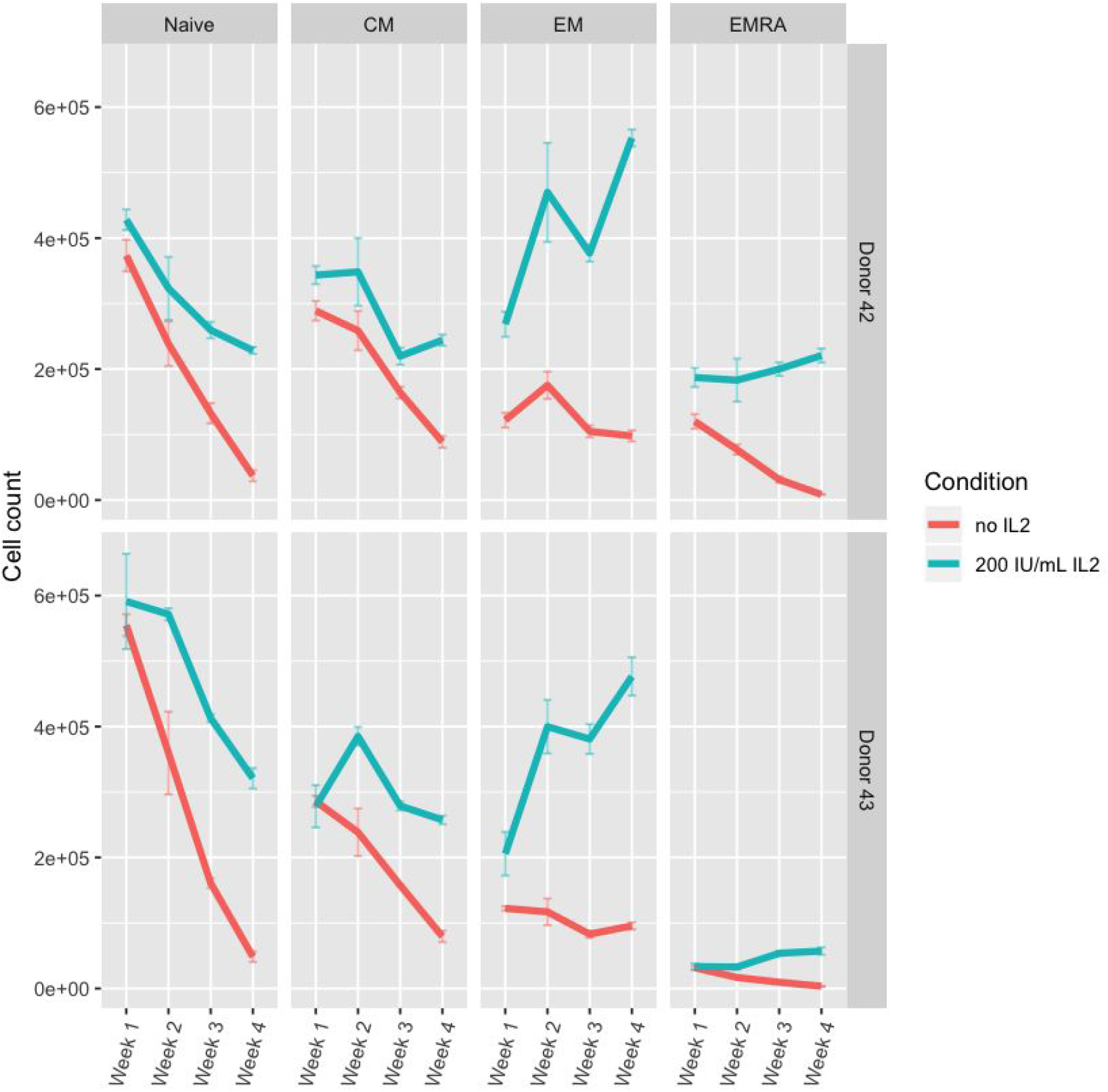
Long-term culturing of unstimulated T cells, with (200 IU/mL) or without IL-2 supplement, compromises the overall viability of the population and differentially changes the subtype prevalence (n=4 for each donor). The viability for both naive (CCR7+CD45RO-) and central memory (CM: CCR7+CD45RO+) populations monotonically decreased over 4 weeks of culturing regardless of the IL-2 supplement, whereas for both effector memory (EM: CCR7-CD45RO+) and effector memory RA (EMRA: CCR7-CD45RO-), the viability changed differently across two different conditions: IL-2 supplement increased the number of both population (source: <u>Long term culturing of unstimulated cells with or without IL2</u>).

**Supplemental Figure 2:**
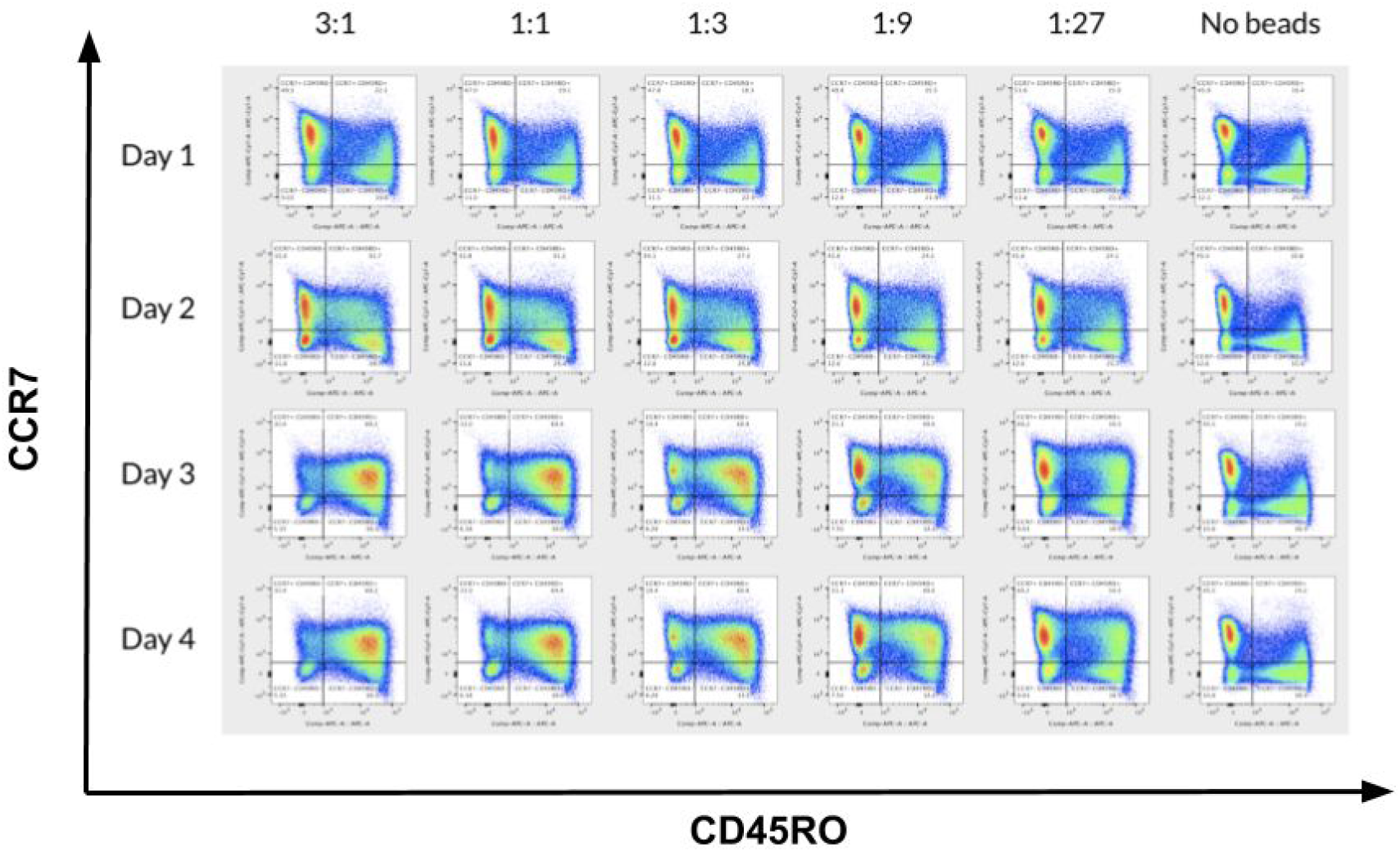
Upon activation with beads, regardless of the bead-to-cell ratio, naive cells (CCR7+CD45RO-) do not fully transition into a central memory phenotype (CM: CCR7+CD45RO+) within the first two days of activation.

**Supplemental Figure 3:**
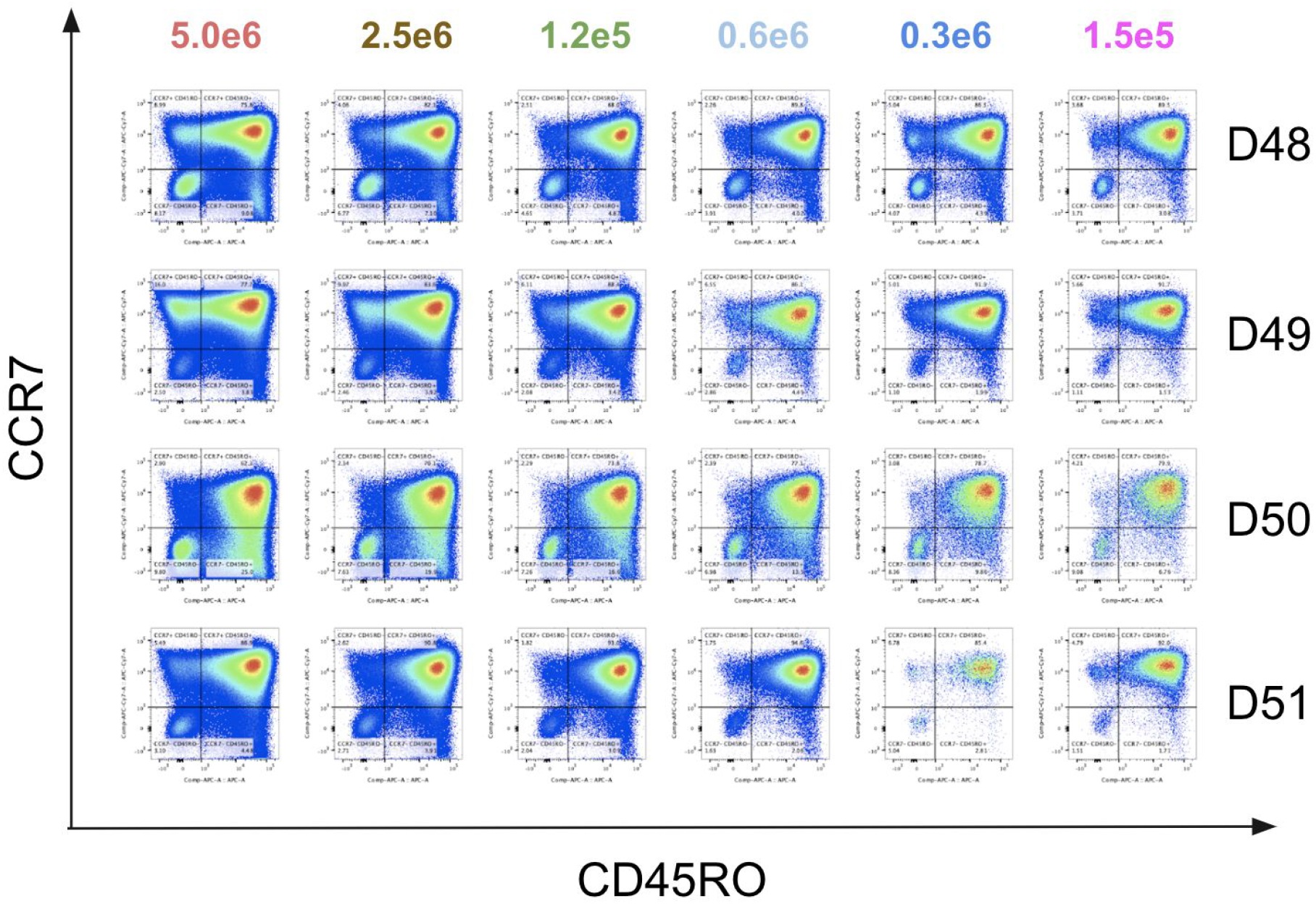
When we activated the T cells varying confluencies (150,000 to 5 million cells per 2 cm^2^ within 1.5 mL of culture media) but using the same bead-to-cell ratio (1-to-1), we saw that confluencies higher than 1.2 million or lower than 300,000 cells per well caused sub-optimal activation. Activation efficiency was estimated based on the fraction of central memory cells (CCR7+CD45RO+) normalized against the fraction of naive cells (CCR7+CD45RO+) across four different donors (D48-D51).

**Supplemental Figure 4:**
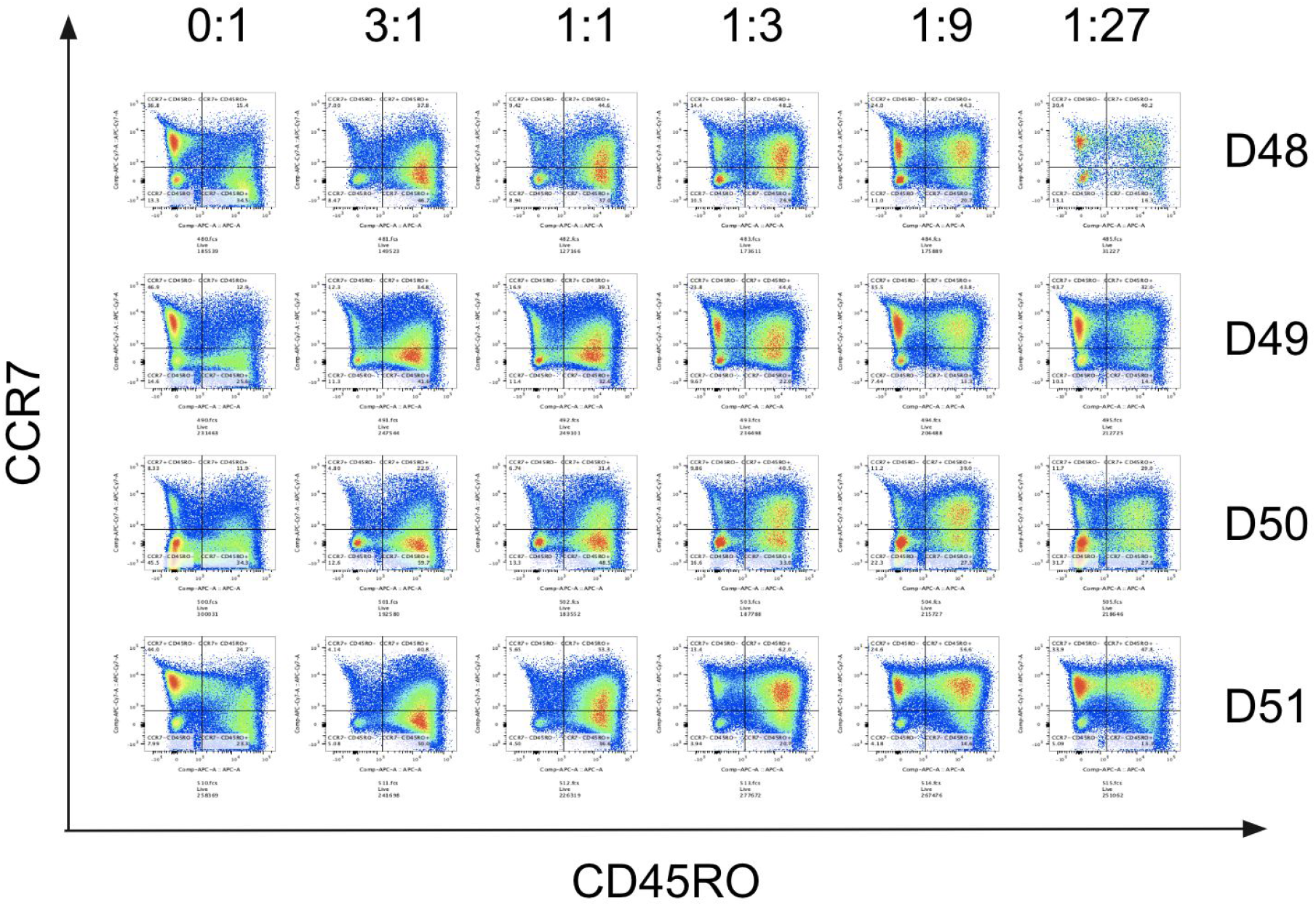
Activating human primary T cells with increasing number of beads leads to decreasing number of naive cells (CCR7+CD45RO-) that are left in the culture upon activation for three days in comparison to no activation 0:1 or no beads) condition across four different donors (D48-D51).

**Supplemental Figure 5:**
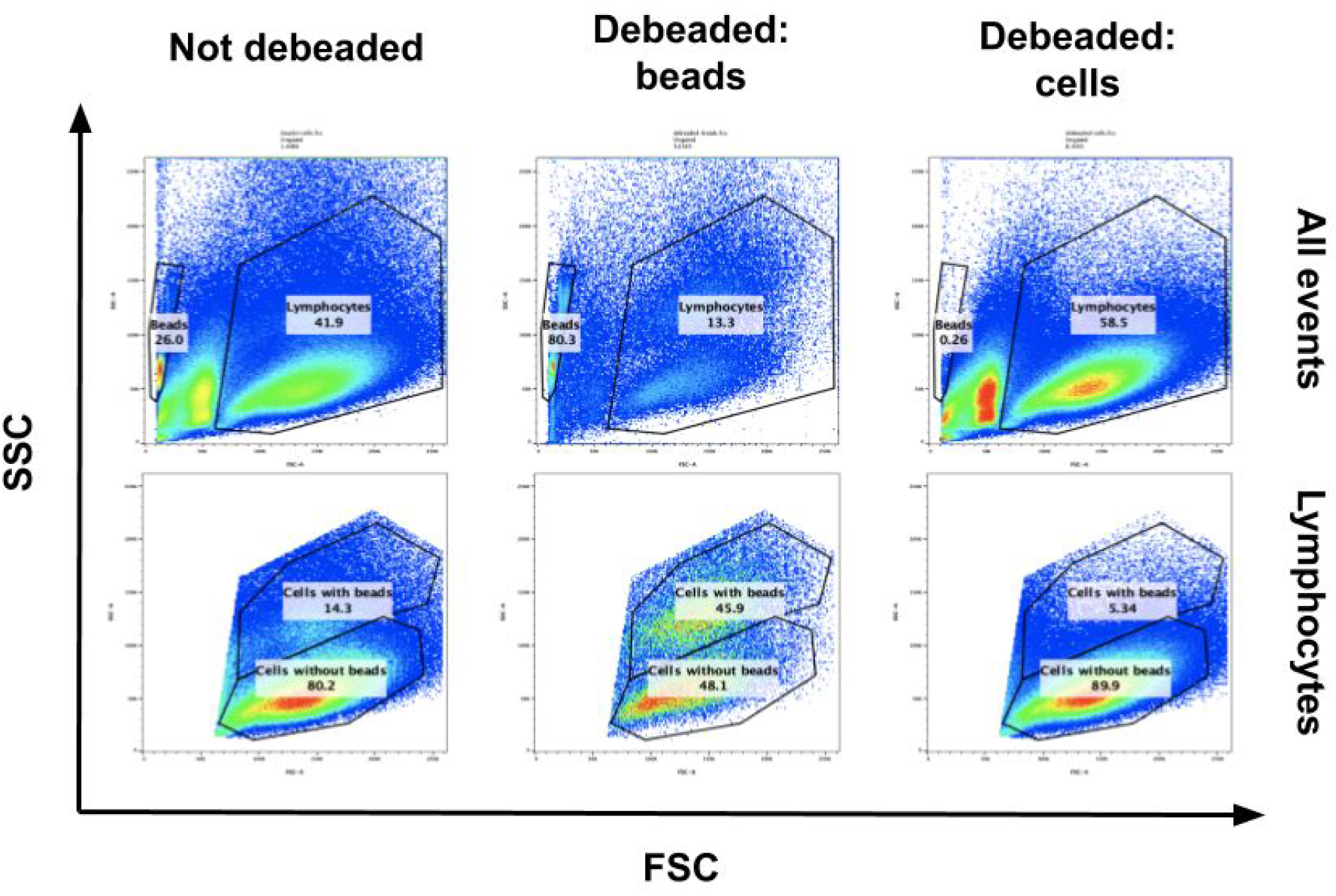
Regardless of cells being debeaded or not, beads (alone) and cells (that are or are not bound to beads) can be distinguished based on their FSC/SSC profiles with the help of flow cytometry. Debeading cells enriches for the lymphocyte population and effectively removed the beads from the culture. Similarly the fraction captured by the magnetic bead during the debeading process was enriched for beads but there were still a considerable amount of cells that had a lymphocyte-like profile. About half of this population had a distinct FSC/SSC profile that was different than the typical debeaded lymphocyte population, assumingly representing cells that were stuck on beads.

**Supplemental Figure 6:**
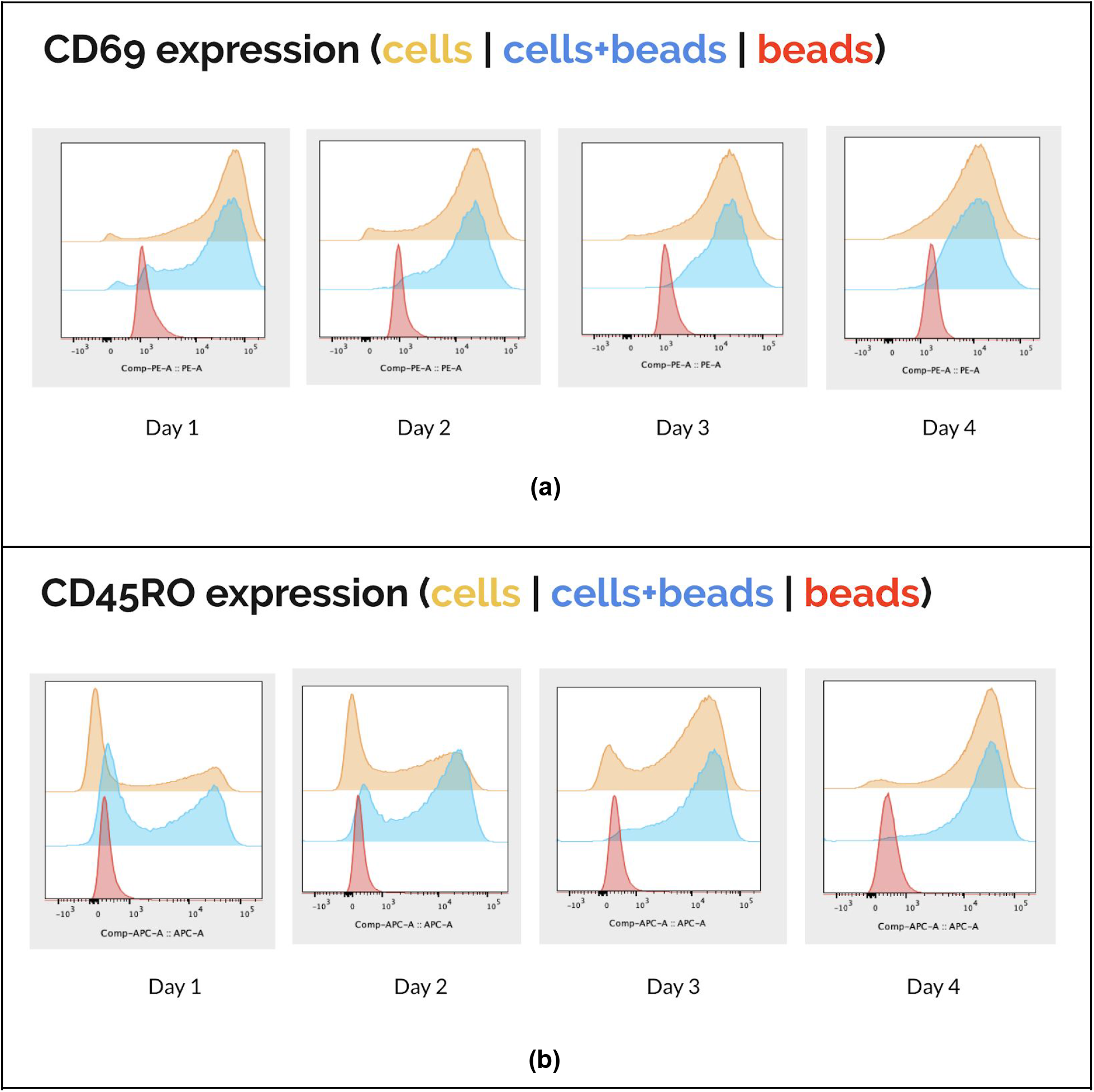
Cells that are stuck on beads (blue) throughout the activation expresses the early activation marker (CD69+; **panel a**) starting from the first day of the activation and predominantly the memory marker (CD45RO+; **panel b**) after the second day of the activation.

**Supplemental Figure 7:**
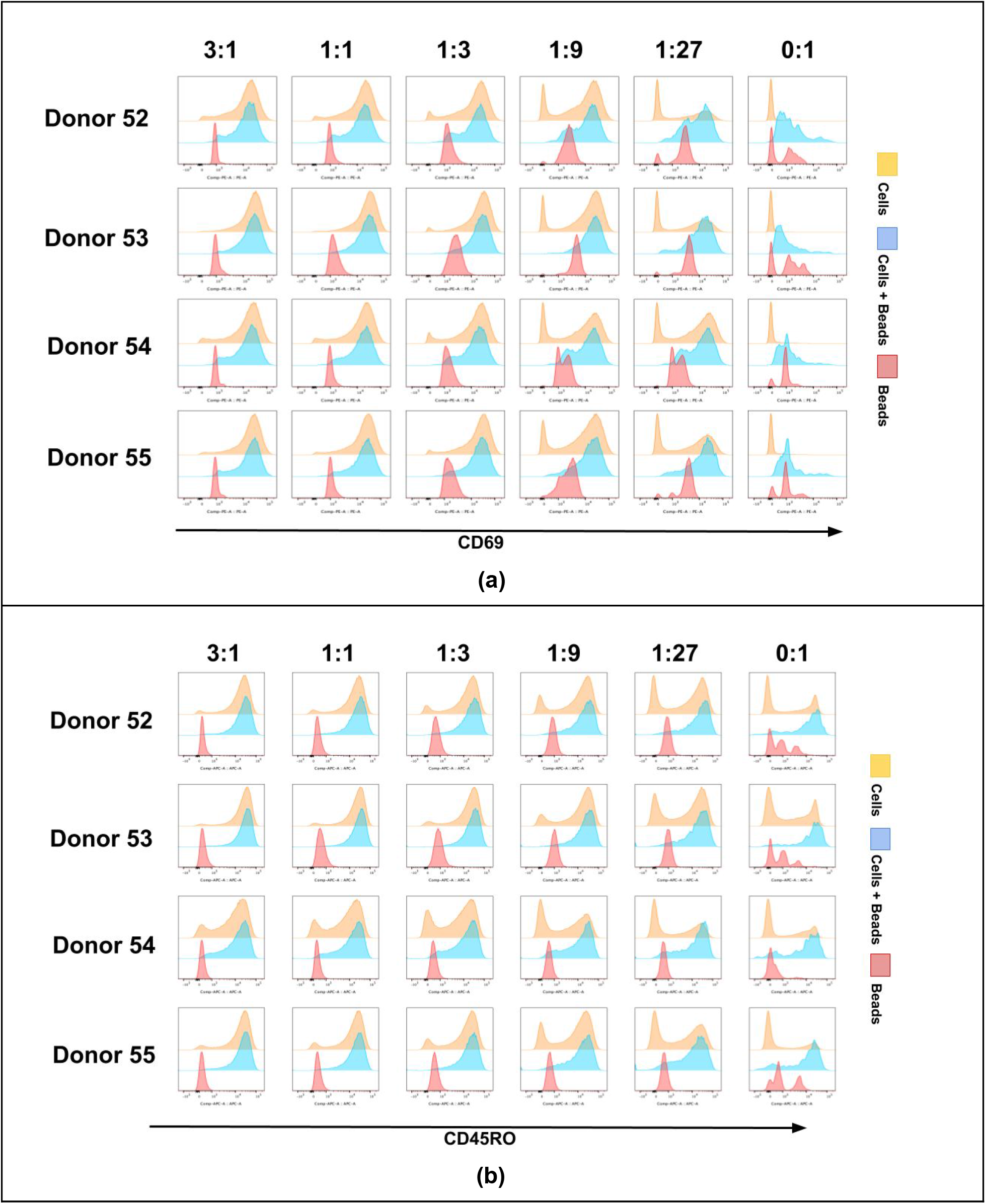
Cells that are stuck on beads (blue) show signs of early activation (CD69+; **panel a**) on the second day of the activation and are of memory phenotype (CD45RO+; **panel b**) by the third day of the activation. Decreasing the number of beads also decreases the overall early-activated and memory-phenotype T cell yield (yellow).

**Supplemental Figure 8:**
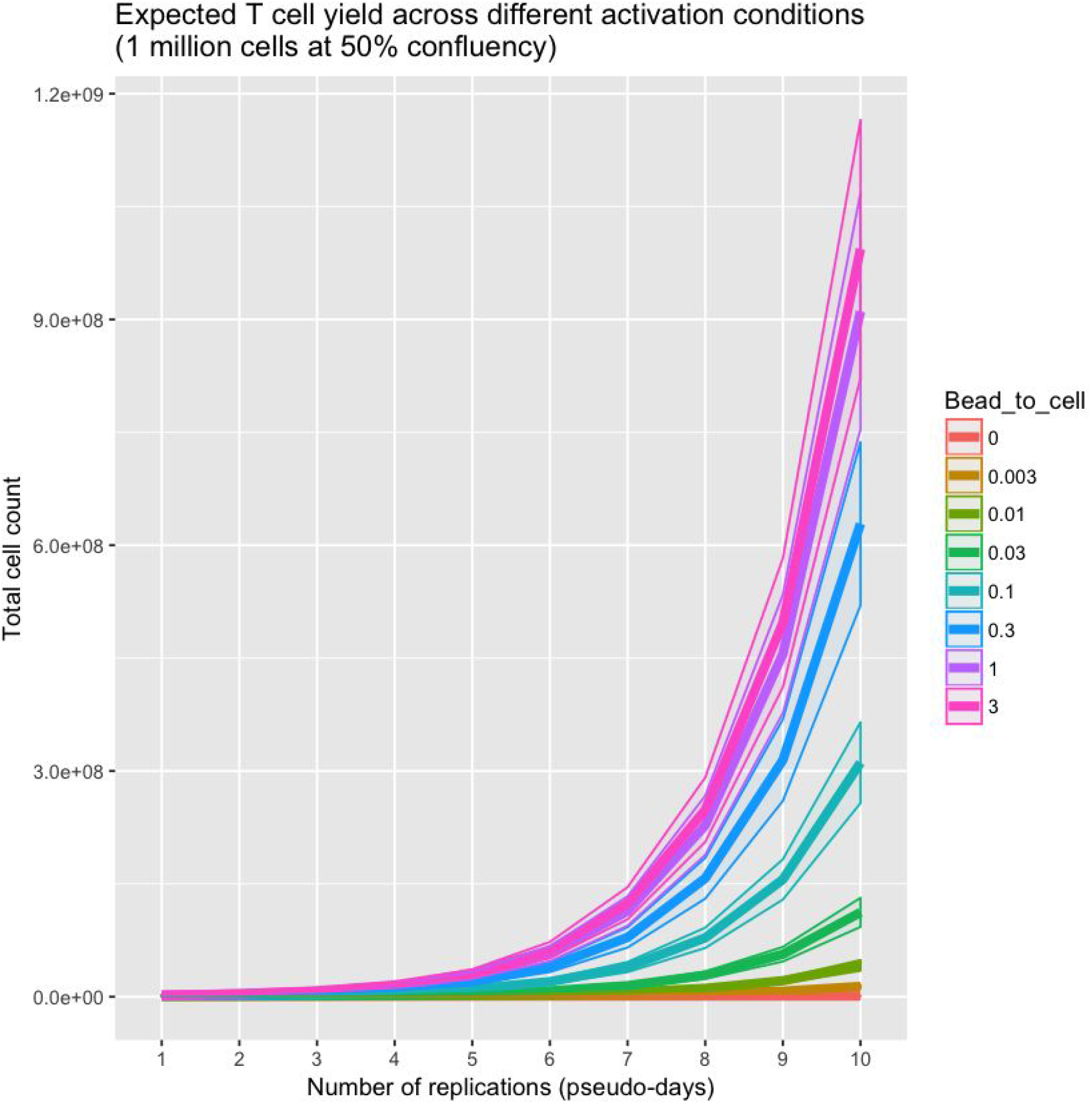
The 95% confidence interval for a typical cell counter is +/- 170,000 cells per 1 million cells per 1 mL, which means that our capabilities of distinguishing different activation efficiencies by considering the final number of cells that are in the culture within the long-term are very limited. Our simulations showed that at a 50% hypothetical confluency, the cell counter could not tell different T cell populations that were activated at different bead-to-cell ratios, namely 3-to-1, 1 -to-1, and 1-to-3 (source: <u>Modeling bead-based T cell activation on a population level</u>).

